# A case-control neuroimaging investigation of chronic Zika virus-infected adults

**DOI:** 10.1101/2025.03.05.641727

**Authors:** Suhnyoung Jun, Richard Bido-Medina, Jairo Oviedo, Isidro Miches, Daniel Llano, Luis Tusen, Peter Stoeter, Minelly Rodriguez, Sepideh Sadaghiani

## Abstract

Systemic viral infections with neurotropic potential pose significant global health challenges. The Zika virus (ZIKV) is known for its pronounced neurotropism, with recent infectious clusters raising renewed public health concerns. While research has predominantly focused on congenital populations, growing evidence suggests that the mature central nervous system (CNS) is also vulnerable. However, no study has examined the long-term impact of ZIKV infection on the adult human brain.

To address this gap, we studied a rare group of adult ZIKV patients presenting with both peripheral (Guillain-Barré Syndrome; GBS) and CNS-related neurological symptoms. We compared these patients at the chronic stage (5 to 12 months post-infection) to healthy controls and to patients with GBS of non-ZIKV etiology (total *N*=43). Structural and functional measures included cortical thickness, white matter hyperintensities, diffusion metrics, and resting-state functional connectivity.

Despite the rarity of both patient populations, power analyses indicated that our sample size could detect large group differences—effect sizes deemed reasonable given the severity of neurological symptoms in the ZIKV group. Nonetheless, our multimodal analyses yielded null results, with Bayesian statistics (where applicable) providing evidence for a lack of effects.

The null findings suggest that chronic ZIKV infection in adults is not associated with brain changes of large magnitude. Importantly, this study offers detailed clinical characterization of a heavily understudied group. In light of recent ZIKV outbreaks, this characterization underscores the need to monitor, study, and provide longitudinal care to adult survivors of severe ZIKV infections.

## 1. Introduction

The Zika virus (ZIKV), a neurotropic flavivirus, gained global attention following its explosive spread across French Polynesia and the Americas in 2015 (Rasmussen et al., 2016). As of December 2022 (most recent WHO report; World Health Organization (WHO), 2022), a total of 89 countries and territories have documented evidence of autochthonous mosquito-borne transmission of ZIKV. Importantly, recent large-scale outbreaks in Asia in 2024 (de Jong and Grobusch, 2025) and in South America–including Argentina and Brazil– in 2025 underscore ongoing public health threat posed by ZIKV. Specifically, over 42,000 cases were reported across the Americas in 2024 alone (The Pan American Health Organization (PAHO), 2025), with thousands more already confirmed in early 2025.

Although ZIKV research has primarily focused on its devastating effects on fetal brain development—particularly congenital microcephaly—there is increasing recognition that the virus also poses serious neurological risks to adults. Among these, Guillain-Barré Syndrome (GBS), an acute immune-mediated neuropathy of the peripheral nervous system, has been widely documented (Cao-Lormeau et al., 2016; Dos Santos et al., 2016; Munoz et al., 2016, 2017; Parra et al., 2016; da Silva et al., 2017).Supporting a causal link, ZIKV RNA has been detected by reverse transcription polymerase chain reaction (RT-PCR) and other serological methods in different bodily fluids (e.g., blood, cerebrospinal fluids (CSF), and urine) of affected individuals (Parra et al., 2016). Notably, 43% of ZIKV-GBS patients in one study developed neurological symptoms concurrently with or shortly after ZIKV infection, suggesting a para-infectious pattern rather than the classical post-infectious GBS profile (Parra et al., 2016; Munoz et al., 2017).

Beyond GBS, ZIKV has been associated with *central* nervous system (CNS) complications, including encephalopathy, encephalitides, meningitis, myelitis, and seizures (Carteaux et al., 2016; Galliez et al., 2016; da Silva et al., 2017). Despite this growing body of clinical observations, neuroimaging evidence of CNS involvement in adults has been limited primarily to isolated case reports aimed at diagnosis or prognosis (Carteaux et al., 2016; Galliez et al., 2016; da Silva et al., 2017; Pradhan et al., 2017; Sebastian et al., 2017). The estimated risk of developing GBS after ZIKV infection is approximately 2.3 per 10,000 (Malkki, 2016), but the true incidence of CNS complications remains unclear, with only a few cases reports describing CNS complications in ZIKV patients (Carteaux et al., 2016; Galliez et al., 2016; da Silva et al., 2017). Further, reports of ZIKV-infected adult patients presenting with both GBS and CNS symptoms (ZIKV-CNS-GBS) are even more exceedingly rare.

Mechanistic evidence from animal studies further underscores the potential for ZIKV to affect the adult CNS. Mouse models have shown that peripheral ZIKV infection—via intraperitoneal (Aliota et al., 2016; Kim et al., 2020), subcutaneous (Dowall et al., 2016), or intravenous (Papa et al., 2017; Leda et al., 2019) routes— can result in viral entry into the brain. ZIKV may cross into the CNS by breaching the blood-CSF barrier or via transcytosis across the blood-brain barrier, with infected pericytes in the choroid plexus facilitating viral entry into the cortex (Kim et al., 2020). Moreover, ZIKV has shown to preferentially target neural progenitor cells over mature neurons (Sutarjono, 2019), rendering hippocampal structures particularly vulnerable to inflammation, astrocytic activation, premature apoptosis, and autophagy (Li et al., 2016; Souza et al., 2016; Devhare et al., 2017; Zhou et al., 2017; Büttner et al., 2019; Figueiredo et al., 2019; Garber et al., 2019; Bobermin et al., 2020). In parallel, indirect mechanisms —such as the induction of cross-reactive autoantibodies and complement activation—have also been implicated in ZIKV-related neuroinflammation (Rivera-Correa et al., 2019; Davies et al., 2022; Jeong et al., 2023). Together, these findings suggest multiple pathways—both direct and immune-mediated—through which ZIKV may contribute to CNS pathology in human adults.

Despite these converging lines of evidence, the extreme rarity of ZIKV-CNS-GBS cases has led to a critical gap in our understanding of this population—particularly in terms of systematic clinical characterization and neuroimaging-based evidence. To our knowledge, no prior case-control neuroimaging study has investigated the impact of ZIKV infection on adult brain structure and function at the chronic stage. Our group previously addressed part of this gap by conducting the first study to evaluate brain alterations in this rare population at the subacute phase of infection, identifying changes in gray matter volume and resting-state functional connectivity in the ZIKV-CNS-GBS group compared to healthy controls (Bido-Medina et al., 2018). Building on our prior work, the present study investigated—for the first time—the chronic-stage effects of ZIKV infection on adult brain structure and function in this same rare cohort.

Specifically, we examined both structural and functional brain integrity in ZIKV-CNS-GBS patients at the chronic stage (5–12 months post-infection). Structural measures included cortical thickness, white matter lesions, and white matter integrity assessed via diffusion-tensor imaging. Functional connectivity analysis focused on the hippocampus, a region known to be particularly susceptible to ZIKV-related damage. Given the rarity of this clinical population, our sample consisted of 14 ZIKV-CNS-GBS patients along with two control groups: 14 healthy controls to establish a normative baseline, and 15 patients with classical GBS of non-ZIKV etiology (nonZIKV-GBS). This design allowed us to better isolate brain alterations specifically attributable to ZIKV-associated CNS involvement, independent of GBS. While the sample size permits detection of large group differences, the severity, rapid onset, and debilitating nature of symptoms in this cohort warrants investigation into potential large-scale brain alterations. By including both healthy and disease control groups, this study aims to clarify the long-term structural and functional effects of ZIKV infection and contribute to a more comprehensive clinical understanding of this rare but clinically significant population.

## 2. Materials and Methods

Figure 1 is a schematic representation of the overall approach and analysis subsections. Our study included a comprehensive set of structural features and employed a consistent analytical approach to investigate the long-term effect of adult ZIKV infection on structural and functional changes.

**Figure 1.**
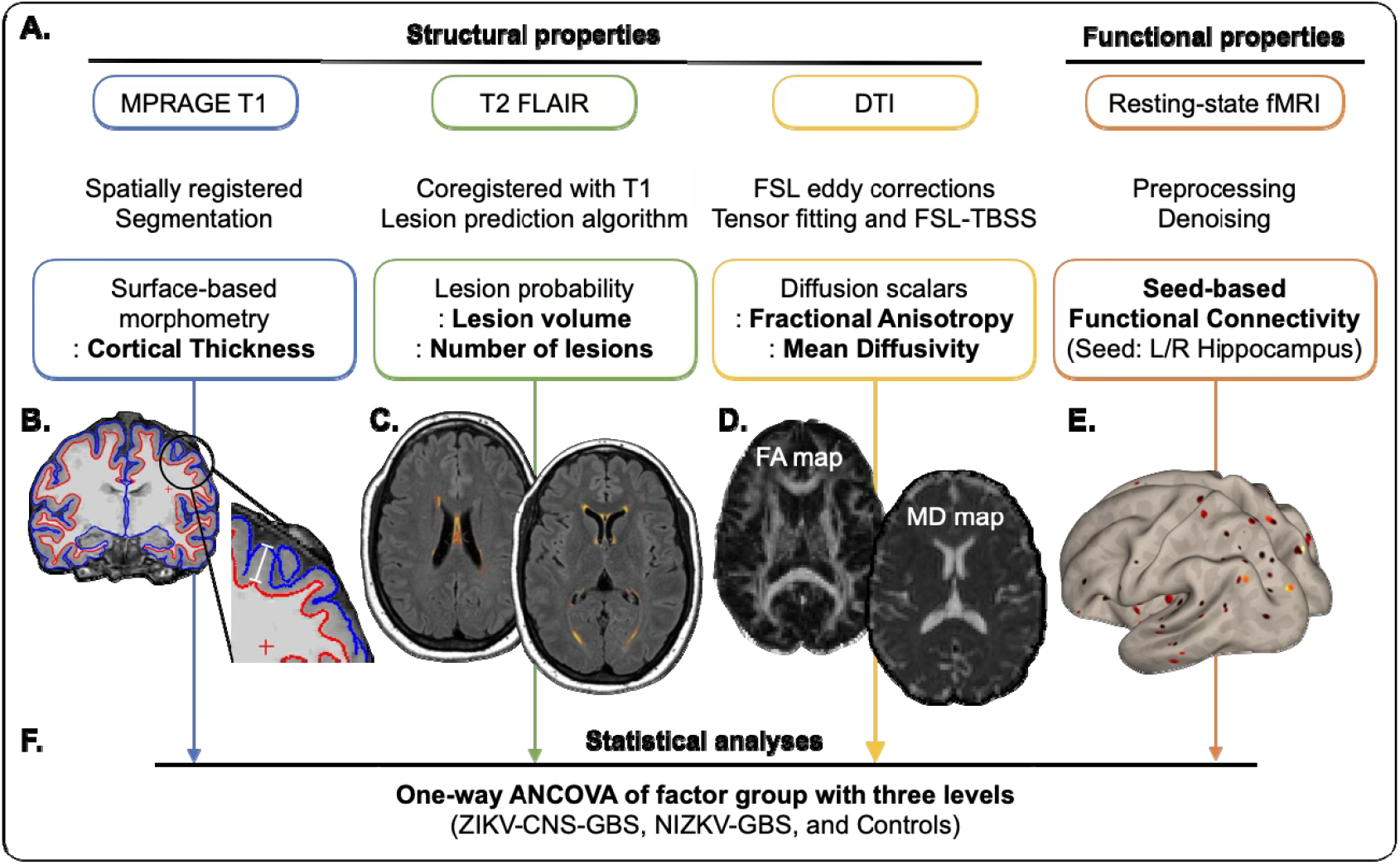
An overview of the analysis pipeline. This figure illustrates the workflow for analyzing multiple neuroimaging modalities. **(A)** For each modality, preprocessing was performed and key structural and functional features were extracted. Specifically, cortical thickness was assessed using surface-based morphometry; lesion characteristics were evaluated via lesion probability mapping; white matter integrity was examined through diffusion metrics—including fractional anisotropy (FA) and mean diffusivity (MD); and resting-state functional connectivity was analyzed using a seed-based approach. **(B–E)** Panels provide illustrative examples to aid understanding of the metrics derived from each modality. **(B)** Cortical thickness was defined as the distance between the white matter surface (red boundary) and the pial surface (blue boundary). **(C)** White matter hyperintensities were identified using an automated algorithm as visualized here in warm colors. **(D)** FA maps reflect the degree of directionality of water diffusion, while MD maps indicate the overall magnitude of water diffusion regardless of direction. Brighter colors represent higher values. **(E)** The functional connectivity map shows the brain regions functionally connected to a predefined seed region (e.g., the left hippocampus), with brighter colors denoting stronger connectivity. **(F)** Each of these features was then subjected to a one-way ANCOVA to examine group differences across three cohorts: adults with confirmed ZIKV-associated CNS and GBS symptoms (ZIKV-CNS-GBS), adults with non-ZIKV-related GBS (nonZIKV-GBS), and healthy controls. The ANCOVA was performed in a voxel-wise manner (except for FLAIR-based measures) and subjected to multiple comparisons corrections.

### 2.1. Participants

We recruited 14 adult patients (6 females; age = 34.93 (± 8.14) years) with confirmed ZIKV infection and ZIKV-associated CNS-related neurological symptoms and GBS manifestations (ZIKV-CNS-GBS), 15 adult patients (5 females; age = 43.21 (±9.37) years) with GBS manifestations without ZIKV etiology (nonZIKV-GBS), and 14 healthy controls (6 females; age = 32.21 (±4.98) years). All subject were recruited in the Dominican Republic, endemic to Aedes mosquito-borne diseases. Recruitment and clinical assessment occurred at Hospital Salvador B. Gautier, in Santo Domingo, and MRI data were acquired at Centro de Diagnóstico Medicina Avanzada y Telemedicina (CEDIMAT), Santo Domingo, Dominican Republic. All subjects gave written informed consent according to procedures approved by the Internal Review Boards of the University of Illinois at Urbana-Champaign, IL, USA, Hospital Gautier, and CEDIMAT. All patients presented an abrupt onset of neurological deterioration with variable periods between systemic manifestation and neurological symptoms. Only patients with stable cognitive and clinical status (Mini-Mental State Examination >= 23 and Glasgow scale >= 13) underwent the MRI scan.

All patients (ZIKV-CNS-GBS and nonZIKV-GBS) were diagnosed with a GBS-like syndrome by clinical manifestations and electromyography (EMG), and CSF testing was performed (i.e., albumin-cytologic dissociation) in 18 of the patients (i.e., 10 ZIKV-CNS-GBS patients and 8 nonZIKV-GBS patients). Nerve velocity conduction using EMG (EMG/ NVC) was assessed by investigator L.T. during the acute stage. HIV status was assessed using ELISA and was reported negative for all patients. Disability due to the radiculopathy was measured by Hughes modified scale at admission and discharge. In accordance with the Dominican Protocol (2016) for the management of GBS, patients were treated with immunoglobulins (2 g/kg divided in 2–5 doses of antibodies unlikely to affect brain function) during the first 5 days of onset of neurological manifestations. Patients did not receive specific medication or treatment at time of scan.

In all patients (ZIKV-CNS-GBS and nonZIKV-GBS) and upon admission to the neurology service, Herpes Simplex 1 (HSV-1) was excluded through immunologic testing (IgM/IgG enzyme-linked immunosorbent assay (ELISA). Other potential febrile syndrome etiologies (Dengue, Chikungunya, Malaria, Leptospirosis, Mononucleosis infecciosa, and HIV primo infection) were also ruled out using immunological tests according to the febrile syndromes protocol of the Dominican Republic. Venereal Disease Research Laboratory (VDRL) testing was performed to exclude spirochete-related CNS diseases (e.g., syphilis, Lyme’s); all resulting non-reactive. Treating neurologists also excluded other conditions with potential CNS involvement, including viral encephalitis, acute disseminated encephalomyelitis (ADEM), and meningitis, based on clinical presentation, cerebrospinal fluid (CSF) analysis, and electromyography (EMG) findings.

In the ZIKV-CNS-GBS, the onset of neurological symptoms occurred within 2 to 30 days (mean = 8.85 days ± 8.03) after the instauration of febrile symptoms associated with ZIKV infection, following a para-infectious pattern with rapid onset. The onset of GBS occurred within 1 to 5 days (mean = 2.43 days ± 1.22) after the above-mentioned onset of other neurological symptoms. Upon admission to the neurology service, recent ZIKV infection was confirmed through immunologic testing (IgM/IgG enzyme-linked immunosorbent assay (ELISA) serum levels > 1.1 mg/dL). The patients later returned for neuroimaging at the chronic stage (12 months ± 10.18 after the initial MRI assessment). CNS and PNS symptoms assessed at this timepoint are reported in Table 1.

**Table 1.** Clinical manifestations of chronic-stage ZIKV-CNS-GBS and nonZIKV-GBS patients in the study.

| Manifestations | ZIKV-CNS-GBS (n = 14) | nonZIKV-GBS (n = 15) |
| --- | --- | --- |
| Guillain–Barre variant (# of patients with atypical descending pattern) | 5 |  |
| Guillain–Barre variant (# of patients with ascending pattern) | 9 | 15 |
| **Central Nervous System (CNS)** |  |  |
| Bitemporal Hemianopsia |  |  |
| Ophthalmoplegia | 1 |  |
| Sialorrhea | 1 |  |
| Hands Dystonia | 1 (bilateral) |  |
| Central facial palsy | 3 | 2 |
| Fatigue | 1 |  |
| Headache | 3 | 4 |
| Loss of visual acuity | 1 | 5 |
| Photophobia | 1 | 1 |
| Peripheral Facial Paresis | 1 (right) |  |
| Short-term Memory Disturbances | 1 |  |
| **Peripheral Nervous System (PNS)** |  |  |
| Calf pain | 1 |  |
| Dysphonia | 1 |  |
| Upper Limb weakness | 2 (1 distal) |  |
| Areflexia | 5 | 14 |
| Dysesthesia | 8 (1 all limbs, 1 right limb) | 7 |
| Loss of sphincter tone | 5 | 10 |
| Muscular atrophy |  | 2 |
| Paraplegia | 2 | 6 |
| Paresthesia | 1 | 5 |
| Tetraparesis | 3 | 10 |

In the nonZIKV-GBS group, the onset of neurological symptoms occurred without evidence of recent ZIKV infection, as demonstrated by negative immunological testing upon admission. The patients later returned for neuroimaging at the chronic stage (10.95 months ± 9.58 after the initial onset of the GBS-associated neurological symptoms), with both CNS and PNS symptoms assessed at the same time (see Table 1).

The healthy control group consisted of individuals with no history of neurological disorders, recent infections, or febrile illnesses.

Values represent the number of patients exhibiting each manifestation within the specified group. Individual patients may present with more than one manifestation. Symptoms were assessed at the chronic stage, concurrent with the neuroimaging scan.

### 2.2. MRI data acquisition

All MRI data were acquired on a 3 Tesla Philips Achieva scanner (Philips Healthcare, Best, The Netherlands) at CEDIMAT facilities. We acquired T1-weighted Sagittal MPRAGE sequence (TR/TE = 3000/2.6 ms, flip angle = 90°, voxel size = 1 × 1 × 1 mm^3^, 180 slices), T2-weighted axial Fluid Attenuation Inversion Recovery (FLAIR) sequence (TR/TE = 11000/120 ms, inversion time (TI) = 2800 ms, flip angle = 120°, voxel size = 1 × 1 × 1 mm^3^, 150 slices), and 6 min eyes closed resting state functional MRI (fMRI) using T2-weighted EPI sequence (TR/TE = 2000/30 ms, in-plane matrix size = 80 × 80, flip angle = 80°, voxel size = 2.38 × 2.4 × 3 mm^3^ without slice gap, 34 axial slices aligned to AC-PC plane). Diffusion tensor imaging (DTI) was acquired using a single shot, diffusion weighted spin-echo EPI sequence (EPI factor = 67, TR/TE = 5900/60 ms, flip angle = 90°, voxel size = 2 × 2 × 2 mm^3^ without slice gap, 33 contiguous slices, in-plane matrix size = 128 × 128, water-fat shift (WFS)/bandwidth (BW) = 20.027 pixels/21.7 Hz, BW in EPI frequency direction = 2502.6 Hz, and total scan time = 4.33 min). Maximum *b*-value was 1000 s/mm^2^ in 32 diffusion-weighted directions, and one volume was acquired without diffusion weighting (*b*_0_ = 0 s/mm^2^).

### 2.3. Tract-Based Spatial Statistics

We processed the diffusion imaging data using FSL (FMRIB Software Library) packages and employed the tract-based spatial statistics (TBSS) method (Smith et al., 2006) to derive diffusion tensor parameters: fractional anisotropy (FA) and mean diffusivity (MD). Briefly, eddy-current and motion corrections were applied, followed by fitting the diffusion tensor to the raw diffusion data to calculate FA and MD maps using default parameters (Smith, 2002; Smith et al., 2006; Jenkinson et al., 2012). The FA images were aligned to a 1×1×1 mm^3^ FMRIB58_FA standard space, and individual FA images were projected onto the mean fiber tract skeleton. Similarly, all MD images were warped to a common space using the non-linear registration created from the FA analysis. The warped MD images of all subjects were merged into a 4-dimensional file and projected onto the original mean FA skeleton. The resulting FA and MD images were then fed into voxel-wise cross-subject statistics.

### 2.4. Preprocessing of functional and anatomical MRI data

The preprocessing of functional MRI (fMRI) data was conducted using *fMRIPrep* (version 22.0.2; (Esteban et al., 2019)), an analysis-agnostic pipeline that minimizes manual intervention and optimizes data quality. Briefly, anatomical preprocessing involved skull-stripping of T1-weighted images using Advanced Normalization Tools (ANTs), followed by segmentation into gray matter, white matter, and cerebrospinal fluid with FSL’s FAST (Zhang et al., 2001), and cortical surface reconstruction via FreeSurfer (Dale et al., 1999; Fischl, 2012). Spatial normalization was performed to MNI152NLin6Asym and MNI152NLin2009cAsym templates using ANTs. Functional preprocessing included generating a BOLD reference image, correcting for head motion using FSL’s MCFLIRT (Jenkinson et al., 2002), and slice-timing correction with AFNI (Cox and Hyde, 1997). Susceptibility distortion correction was applied using either field maps or a fieldmap-less method based on T1-weighted image registration, followed by alignment of BOLD images to T1-weighted data with boundary-based registration (Greve and Fischl, 2009). The functional data were resampled to both native and standard MNI spaces with minimal smoothing.

### 2.5. Surface-based morphometry

Surface-based morphometry (SBM) was conducted to derive whole-brain morphometric measures using the default preprocessing steps of the Computational Anatomy Toolbox (CAT12 version 12.8.1) implemented in SPM12 (version 7771). Briefly, the high-resolution MPRAGE T1 data were segmented, and brain surface meshes were registered to the ‘FsAverage’ template in FreeSurfer. After topology correction, spherical mapping and spherical registration were conducted using the DARTEL algorithm. The Projection-based Thickness (PBT) method was then used to estimate Cortical Thickness by calculating the distance between the gray matter and white matter boundaries (Dahnke et al., 2013). Unlike voxel-based morphometry (VBM), which provides a composite measure of gray matter density and volume, PBT allows for the separation of Cortical Thickness, surface area, and folding. This reduces the partial volume effects inherent to VBM and provides more accurate and sensitive measurements of localized changes in cortical structures (Hutton et al., 2009; Riccelli et al., 2017). The Cortical Thickness maps for the left and right hemispheres were resampled into template space and smoothed using a 15-mm full-width at half-maximum (FWHM) Gaussian kernel.

### 2.6. Lesion segmentation

Cerebral white matter lesions appear as hyperintensities on FLAIR images. We used a semi-automatic lesion prediction algorithm (LPA) (Schmidt, 2017), implemented in the SPM12-based Lesion Segmentation Toolbox (LST version 3.0.0), that provides automatic lesion filling on the MPRAGE T1-coregistered FLAIR images without needing parameter optimization or binary thresholding of the lesion masks using the logistic regression approach. The resulting lesion probability maps provided two main parameters: Total Lesion Volume (ml) and a Total Number of Lesions.

### 2.7. Functional connectivity analysis

Seed-based functional connectivity analysis was conducted with a focus on the hippocampus as a seed due to its known susceptibility to the ZIKV (see introduction). We used the *CONN* toolbox (release 22.v2407) and SPM release 12.7771. The *fMRIPrep*-processed functional data, along with FreeSurfer-segmented anatomical data, were imported into *CONN*. A Gaussian kernel of 6mm with full width at half maximum (FWHM) was used to smooth the functional data for coherence enhancement. Seed-to-voxel correlations were computed by extracting average time series from the region-of-interests (ROIs) and correlating them with all brain voxels, with Fisher z-transformation applied to improve normality. Specifically, the ROIs were defined as bilateral hippocampal areas using the Automated Anatomical Atlas 3 (AAL3; (Rolls et al., 2020)). These ROIs were chosen based on the hypothesis that the progenitor-targeting ZIKV will affect hippocampal area where adult neurogenesis occurs in human brain (Li et al., 2016; Moreno-Jiménez et al., 2021). Significance was assessed after cluster-based correction for multiple comparisons over voxels.

Exploratory whole-brain functional connectivity analysis was additionally performed to assess any potential effects beyond hippocampal connectivity. Specifically, a functional connectivity matrix (148 × 148) was computed for each subject using the Destrieux atlas with 148 regions (Destrieux et al., 2010). To identify group differences in large-scale network connectivity, we employed the network-based statistics (NBS) approach (Zalesky et al., 2010). In connectivity analyses, the multiple comparisons problem arises due to the large number of statistical tests performed across connections, increasing the risk of false positives. NBS addresses this by identifying clusters of connections that show significant between-group differences and evaluating the statistical significance of these clusters as a whole, while controlling for the family-wise error rate (FWER).

### 2.8. Statistical analysis

For each diffusion parameter (FA and MD), we conducted a non-parametric one-way ANCOVA, adjusting for age and sex, using the threshold-free cluster enhancement (TFCE) approach (Smith and Nichols, 2009). The TFCE method involved 5000 randomized permutations, with a family-wise error (FWE) corrected significance threshold of *p* < 0.05. This produced maps of multiple comparisons-corrected, cluster-enhanced statistics across space. Cluster coordinates and white matter tract labels were identified using the JHU-ICBM-DTI-81 WM atlas.

We investigated group differences in Cortical Thickness using a one-way ANCOVA with three levels (ZIKV-CNS-GBS, nonZIKV-GBS, and healthy controls), adjusting for age and sex. This analysis was implemented using SPM12. Statistical significance was determined with a voxel-level threshold of FWE *p* < 0.05 and a cluster-level threshold of uncorrected *p* < 0.05.

Group differences in white matter lesion parameters (Total Lesion Volume and Total Number of Lesions) were tested using a one-way ANCOVA with three levels, adjusted for age, sex, and TIV, implemented using JASP (version 0.16.2).

For each seed (left and right hippocampus), we conducted a one-way ANCOVA with three levels, adjusting for age, sex, and head motion (mean framewise displacement (FD); Power et al., 2014). Statistical significance was determined with a voxel-level threshold of FWE *p* < 0.05 and a cluster-level threshold of uncorrected *p* < 0.05. Note that the one-way ANCOVA of the factor group, adjusted for age and sex, on the mean FD (Power et al., 2014) revealed no significant difference in head motion among the groups: *F*(_2, 31_) = 1.10, *P* = 0.345.

## 3. Results

The neurological symptoms in the ZIKV-CNS-GBS differed from nonZIKV-GBS patients in terms of the rapid peri-infectious onset, the high prevalence of CNS symptoms, and atypical descending pattern of GBS progression in a subset of cases (Table 1). Below, we detail multi-modal comparisons between the two patient groups as well as healthy controls.

### 3.1. Lack of Evidence for Adult ZIKV-CNS-associated Changes in White Matter Hyperintensities

White matter regions with hyperintensities were identified using an automated FLAIR-based LPA method. Due to missing FLAIR scans, one ZIKA-CNS-GBS patient and one healthy control subject were excluded, resulting in a total of 41 subjects for statistical analysis. Visual inspection of the white matter lesion maps revealed that most hyperintensities were located in the periventricular regions. However, there was no evidence to support the presence of ZIKV-associated white matter lesions. A one-way ANCOVA for Total Lesion Volume 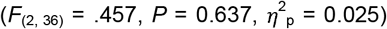 and Total Number of Lesions 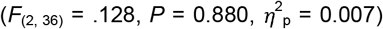 showed no significant group differences. With 41 subjects across three groups and three covariates (i.e., age, sex, and TIV), this ANCOVA had 80% power to detect a large effect size *f* = 0.51 at an uncorrected error probability of 0.05. Therefore, the null result suggests that a large effect is unlikely to be present.

To directly assess the likelihood of the null hypothesis, we calculated Bayes Factor (BF) values (H_0_: no group effect) versus the alternative hypothesis (H_1_). The Bayes Factor for Total Lesion Volume (BF_01_ = 3.94) and Total Number of Lesions (BF_01_ = 4.57) provided moderate evidence in favor of H_0_, suggesting that the data are more likely to occur under the null hypothesis than under the alternative. In contrast and as expected, there was a strong effect of age on white matter lesions: Total Lesion Volume 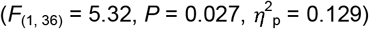 and Total Number of Lesions 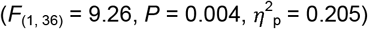. Full information on the Total Number of Lesions and Total Lesion Volume of the subjects is reported in *Table S1*.

### 3.2. Lack of Evidence for Adult ZIKV-CNS-associated Changes in White Matter Microstructure

We assessed white matter microstructural properties using two diffusion-tensor scalars, FA and MD. FA measures directional diffusion, reflecting white matter cohesion, while MD provides a directionality-independent measure of diffusivity, sensitive to changes in myelin and intra/extracellular space (Vuoksimaa et al., 2016). Lower FA indicates reduced anisotropic diffusion, while lower MD suggests more restriction to diffusion across all directions. A one-way ANCOVA was conducted for each scalar to examine group differences, corrected for age and sex (TFCE-corrected *P (P*_*TFCE*_) < .05) and no group differences were observed in either measure. Note that with 42 subjects across three groups and two covariates (i.e., age and sex), this ANCOVA has 80% power to detect a large effect size (*f* = 0.50 at an uncorrected error probability of 0.05). Therefore, the null findings suggest that group effects, if they exist, do not reach a large effect size.

### 3.3. Lack of Evidence for Adult ZIKV-CNS-associated Changes in Cortical Thickness

The one-way ANCOVA of the factor group on Cortical Thickness did not reveal significant differences among the groups. With 43 subjects across three groups and two covariates (i.e., age and sex), this ANCOVA had 80% power to detect a large effect size *f* = 0.49 at an uncorrected error probability of 0.05. The null result thus indicates the absence of at least a large effect.

### 3.4. Lack of Evidence for Adult ZIKV-CNS-associated Changes in Functional Connectivity

Due to missing functional data from five nonZIKV-GBS patients and two ZIKV-CNS-GBS patients, 36 subjects were included in the analysis. For both seeds, the one-way ANCOVA of the factor group on seed-based functional connectivity did not reveal significant differences among the groups. With 36 subjects across three groups and three covariates (i.e., age, sex, and mean FD), this ANCOVA has 80% power to detect a large effect size *f* = 0.54 at voxel level before cluster-level assessment. The null result therefore shows the absence of large differences across groups. Similarly, the exploratory NBS analysis also resulted in null findings, suggesting that no large-scale alterations in whole-brain functional connectivity were detectable between the groups.

## 4. Discussion

The neurological consequences of ZIKV infection have been overlooked in the adult population. While extensive research on ZIKV has focused on fetal and pediatric outcomes, little is known about the sequalae in adults. his gap is particularly pressing given the absence of approved vaccines or antiviral treatments, coupled with the growing risk of future outbreaks driven by climate change and the broad host range of ZIKV—including mosquitoes, animals, and humans (Vorou, 2016; Munoz et al., 2017; Campos et al., 2020; Baldwin et al., 2021). In this context, our study presents the first case-control investigation into the potential chronic effects of ZIKV infection on adult brain structure and function, using a multimodal neuroimaging approach that included measures of cortical thickness, white matter lesions, diffusion metrics, and resting-state functional connectivity. Although no significant group differences were detected, our findings contribute valuable data to a severely understudied population and underscore the importance of expanding ZIKV research to include long-term adult outcomes.

Notably, the ZIKV-CNS-GBS patients in our study exhibited atypical GBS features, including a rapid onset of neurological symptoms following ZIKV infection and, in some cases, an uncommon descending pattern of symptom progression. This clinical profile is consistent with prior reports indicating a shortened latency period for GBS onset in ZIKV-associated cases—typically 4 to 10 days post-infection (Cao-Lormeau et al., 2016; Parra et al., 2016; Chang et al., 2018; Uncini et al., 2018)—compared to the classical 2–4 week window (Willison et al., 2016) observed in post-infectious GBS of other etiologies. Additionally, ZIKV-related GBS has been associated with non-classical presentations, including regional or asymmetric weakness rather than the typical ascending progression from lower to upper limbs (Kassavetis et al., 2016). These clinical observations provide further support for the distinct neurological profile of ZIKV-CNS-GBS cases and justify their focused investigation.

Although our study did not reveal significant group differences in diffusion measures (FA and MD), which reflect white matter microstructural integrity, prior work has shed light on biological mechanisms through which ZIKV may impact white matter. *In vitro* studies using mature murine CNS cultures have shown that ZIKV can directly damage myelinated fibers and induce axonal pathology, primarily through infection of oligodendrocytes, the myelin-producing cells in the CNS (Schultz et al., 2021b). ZIKV infection has also been associated with enhanced microgliosis and elevated pro-inflammatory cytokine production, including TNF-α (Figueiredo et al., 2019). These inflammatory responses, involving both microglia and astrocytes, can disrupt the formation and maintenance of myelin in white matter tracts (Schultz et al., 2021a). Additionally, both *in vitro* and *in vivo* studies have demonstrated that neurotropic ZIKV induces oxidative stress, leading to mitochondrial dysfunction, DNA damage, and reactive gliosis—processes that may compromise white matter integrity even in the absence of overt structural abnormalities (Almeida et al., 2020; Ledur et al., 2020).

These converging lines of evidence suggest that ZIKV may impair myelinated structures through both direct viral effects and secondary immune-inflammatory mechanisms. Consistent with this, our prior work identified atypical GBS features in ZIKV-infected patients (Bido-Medina et al., 2018), and such non-classical GBS presentations are frequently associated with distinct pathophysiological mechanisms, including the presence of anti-myelin glycolipid antibodies (Rivera-Correa et al., 2019; Davies et al., 2022). These findings support the possibility of a para-infectious humoral response contributing to myelin disruption in ZIKV-CNS-GBS cases, even in the absence of prominent white matter abnormalities detectable through standard imaging metrics.

While our previous study of subacute-phase ZIKV-CNS-GBS patients identified volumetric alterations in gray matter (Bido-Medina et al., 2018), the current investigation—focused on the chronic phase (see *Table S1*) did not reveal significant changes in cortical thickness. This contrast raises the possibility that the gray matter alterations observed during the subacute phase reflected transient neuroinflammatory processes that may have resolved over time. Clinical reports have documented that ZIKV infection in adults can lead to neurological complications such as encephalitis and meningitis (da Silva et al., 2017), which are commonly associated with acute inflammation in the brain. However, by the chronic stage, such inflammation may have subsided or stabilized, resulting in no measurable long-term changes in cortical thickness. Still, the absence of significant group differences in our chronic-phase sample should be interpreted with caution. Given the rarity of this patient population, the sample size was sufficient to detect only large effect sizes; smaller or more localized cortical differences may have been undetected.

The primary limitation of this study arises from the extreme rarity of adult ZIKV patients presenting with both CNS and GBS manifestations, inevitably leading to a small sample size. Efforts to enhance the specificity of our findings by including a disease control group with non-ZIKV GBS further complicated recruitment, given that classical GBS is itself an uncommon condition. As a result, the study was powered to detect only large effect sizes, and smaller or more nuanced group differences may not have been captured.

Despite these limitations, our findings provide the first case-control neuroimaging investigation of chronic-phase ZIKV infection in adults with combined CNS and GBS symptoms. While we did not identify significant group differences with large effect sizes in brain structure or function, this null result does not diminish the clinical significance of the severe neurological symptoms reported in the current study. Instead, the careful characterization of this rare cohort offers a valuable contribution to the limited body of research on adult ZIKV outcomes. It remains possible that subtle or regionally specific alterations exist but were not detected, either due to limited statistical power or because such changes fall outside the broad scope—yet necessarily limited—of the multimodal neuroimaging features assessed in this study. As climate change expands the geographic range of mosquito vectors and recent outbreaks reaffirm the ongoing threat of ZIKV, continued research into adult neurological complications is both timely and essential. Our study underscores the need to monitor adult survivors of severe ZIKV infections and highlights the importance of sustained surveillance and longitudinal care. Expanding the focus of ZIKV research beyond congenital outcomes will be critical for advancing understanding of ZIKV-related neuropathology across the lifespan.

## Acknowledgements

This work was funded by the National Institute of Neurological Disorders and Stroke (NIN/NINDS; R21NS104603).

## Supplementary Materials

**Table S1.** White matter lesions identified with LPA-LST.

| ZIKV-CNS-GBS |  |  | nonZIKV-GBS |  |  | healthy controls |  |  |
| --- | --- | --- | --- | --- | --- | --- | --- | --- |
| Cases | TLV (ml) | TNL | Cases | TLV (ml) | TNL | Cases | TLV (ml) | TNL |
| # 1 | 0.659 | 12 | # 1 | 0.835 | 16 | # 1 | 0.478 | 7 |
| # 2 | 1.012 | 15 | # 2 | 0.773 | 19 | # 2 | 0.599 | 13 |
| # 3 | 0.919 | 15 | # 3 | 0.769 | 10 | # 3 | 0.036 | 2 |
| # 4 | 0.268 | 9 | # 4 | 1.628 | 20 | # 4 | 5.385 | 42 |
| # 5 | 0.643 | 13 | # 5 | 0.786 | 12 | # 5 | 0.235 | 5 |
| # 6 | 0.879 | 17 | # 6 | 1.012 | 22 | # 6 | 0.763 | 9 |
| # 7 | 0.466 | 7 | # 7 | 0.567 | 10 | # 7 | 1.011 | 9 |
| # 8 | 0.621 | 11 | # 8 | 1.582 | 12 | # 8 | 0.649 | 11 |
| # 9 | 0.661 | 8 | # 9 | 0.911 | 16 | # 9 | 2.717 | 16 |
| # 10 | 0.211 | 9 | # 10 | 9.307 | 48 | # 10 | 0.821 | 14 |
| # 11 | 0.088 | 3 | # 11 | 0.573 | 9 | # 11 | 0.266 | 2 |
| # 12 | 1.122 | 12 | # 12 | 1.029 | 16 | # 12 | 0.407 | 13 |
| # 13 | 1.061 | 14 | # 13 | 0.126 | 5 | # 13 | 0.183 | 6 |
|  |  |  | # 14 | 0.831 | 14 |  |  |  |
|  |  |  | # 15 | 1.851 | 18 |  |  |  |
| N cases | Mean (± SD) | Mean (± SD) | N cases | Mean (± SD) | Mean (± SD) | N cases | Mean (± SD) | Mean (± SD) |
| N = 13 | .66 (± .33) | 11.15 (± 3.87) | N = 15 | 1.55 (± 2.28) | 16.50 (± 10.24) | N = 13 | 1.04 (± 1.47) | 11.46 (± 10.20) |

Note: LPA-LST, Lesion Prediction Algorithm in Lesion Segmentation Toolbox; N cases, Number of cases with FLAIR imaging data; SD, standard deviation; TLV, total white matter lesion volume; TNL, total number of white matter lesions; m.d.; missing data.

## Notes

https://doi.org/10.13012/B2IDB-5584114_V1

### Competing Interest Statement

The authors have declared no competing interest.

### Summary of Updates

We are now submitting the null results.

